# Same same but different – The global response of *Escherichia coli* to five different LpxC inhibitors

**DOI:** 10.1101/2023.07.25.550486

**Authors:** Anna-Maria Möller, Melissa Vázquez-Hernández, Blanka Kutscher, Raffael Brysch, Simon Brückner, Emily Marino, Julia Kleetz, Christoph HR Senges, Sina Schäkermann, Julia E Bandow, Franz Narberhaus

## Abstract

A promising but yet clinically unexploited antibiotic target in difficult-to-treat Gram-negative bacteria is LpxC, the key enzyme in the biosynthesis of lipopolysaccharides (LPS), which are the major constituents of the outer membrane. To gain insights into the mode of action of five different LpxC inhibitors, we conducted a comparative phenotypic and proteomic analysis. All five compounds bound to purified LpxC from *Escherichia coli*. Treatment of *E. coli* with these compounds changed the cell shape and stabilized LpxC suggesting that the FtsH-mediated turnover is impaired. LpxC inhibition sensitized *E. coli* to the cell wall antibiotic vancomycin, which typically does not cross the outer membrane. Four of the five compounds led to an accumulation of lyso-PE, a cleavage product of phosphatidylethanolamine (PE), generated by the phospholipase PldA. The combined results suggested an imbalance in phospholipid (PL) and LPS biosynthesis, which was corroborated by the global proteome response to treatment with the LpxC inhibitors. Apart from LpxC itself, FabA and FabB responsible for the biosynthesis of unsaturated fatty acids, were consistently upregulated. Our work also shows that antibiotics targeting the same enzyme do not necessarily elicit identical cellular responses. Compound-specific marker proteins belonged to different functional categories, like stress responses, nucleotide or amino acid metabolism and quorum sensing. These findings provide new insights into common and distinct cellular defense mechanisms against LpxC inhibition. Moreover, they support a delicate balance between LPS and PL biosynthesis with great potential as point of attack for antimicrobial intervention.

**Importance:** The alarming spread of antimicrobial resistance among Gram-negative bacteria calls for novel intervention strategies. Inhibitors of LpxC, the first committed enzyme of lipopolysaccharide biosynthesis have been recognized as promising broad-spectrum antibiotics against Gram-negative pathogens. Despite the development of dozens of chemically diverse LpxC inhibitor molecules, it is essentially unknown how bacteria counteract LpxC inhibition. Our study provides comprehensive insights into the bacterial defense strategies against five different LpxC inhibitors. We show that the cellular response of *Escherichia coli* is compound-specific but shares a common pattern. Inhibition of LpxC is toxic, disrupts membrane integrity, and elicits a stress response, including upregulation of fatty acid biosynthesis proteins. Pre-treatment of *E. coli* with low doses of LpxC inhibitors increased the sensitivity to the cell wall antibiotic vancomycin suggesting new directions in combination therapies.

## 1. Introduction

The ever-increasing antimicrobial resistance threatens public health on a global scale (1, 2). Hence, there is an urgent need for the discovery and development of new antibiotics with novel modes of action. Over the past decades, the looming crisis of multidrug-resistant bacterial pathogens has become acute due to the shortage of novel antibiotics entering the market since the 1970s. Aside from few new classes like oxazolidines and lipopeptides, only derivatives of already existing compounds have been approved by the FDA (3, 4). Classical targets of broad-spectrum antibiotics are essential processes like the bacterial cell wall or protein synthesis. Apart from these, also the membrane biosynthesis has high potential for antibiotic development.

The cell envelope of Gram-negative bacteria like *Escherichia coli* is comprised of a thin peptidoglycan layer flanked by two membranes, the phospholipid (PL)-containing inner membrane and the asymmetrical outer membrane (OM) with PL in the inner leaflet and primarily lipopolysaccharides (LPS) in the outer leaflet. Enzymes involved in LPS biosynthesis are promising antibiotic targets since LPS molecules cover up to 75% of the cell surface and are essential for the maintenance of membrane integrity (5, 6). Importantly, LPS biosynthesis is closely linked to PL biosynthesis, as they share β-hydroxymyristoyl-ACP as their common precursor (7, 8) (Fig. 1A). The acyl-ACP is either converted by one of the two dehydratases FabZ or FabA, enzymes of the fatty acid elongation cycle, or by LpxA, the first enzyme of the lipid A pathway. Lipid A is the highly conserved, membrane-anchored part of LPS. It is connected via a core oligosaccharide to a long sugar chain, the *O*-antigen, whose composition varies between bacterial species (9). As lipid A induces the inflammatory reaction upon host infection, it is a virulence-associated factor often referred to as endotoxin. Lipid A biosynthesis requires nine unique enzymes (9–11). Instead of the first enzyme LpxA, which catalyzes a reversible reaction, the second enzyme, LpxC, is the main driver in this process as it catalyzes the first committed step (Fig.1A) (5). The cellular LpxC level is tightly controlled by posttranslational degradation via the FtsH protease because both too high and too low levels can lead to cell death (12–15). An imbalance in the LPS and PL ratio, e.g., by deletion of *ftsH* or by mutations in *lpxC* can only be tolerated by an adapted rate of PL synthesis (16, 17). These and other findings emphasize the intricate crosstalk between LPS and PL biosynthesis (18, 19), which may be an Achilles heel of diderm bacteria. Due to its role as key enzyme, its high conservation among Gram-negative bacteria, and the lack of homology to mammalian proteins, LpxC is considered the most suitable antibiotic target in the LPS biosynthesis pathway. LpxC is a Zn^2+^-dependent deacetylase, and its level is regulated in a growth phase-related manner to meet the cellular lipid A demand (12, 15, 20). It interacts with its natural substrate, UDP-3-O-(R-3-hydroxymyristoyl)-GlcNAc, at all four segments: uridine, pyrophosphate, glucosamine, and myristate (21).

**FIGURE 1.**
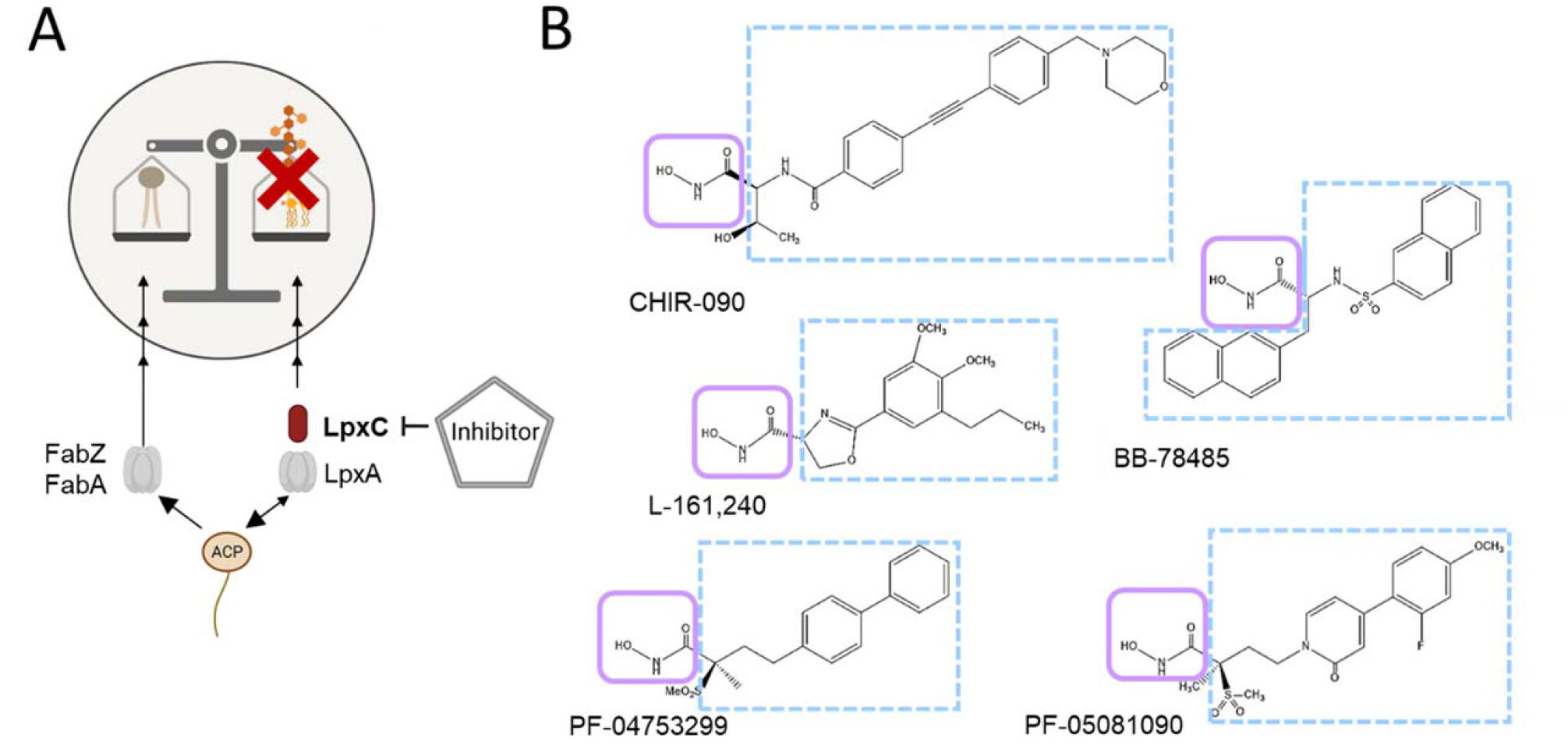
LpxC is a promising antibiotic target due to its critical role in LPS biosynthesis. A) PL and LPS synthesis are linked by the common precursor acyl-ACP. The essential enzyme LpxC catalyzes the first committed step of LPS synthesis and thus represents a promising antibiotic target. The inhibition of LpxC blocks LPS production and critically disrupts membrane integrity. B) In this study, the cellular response to five LpxC inhibitors, the N-aroyl-L-threonine derivative CHIR-090, the aryl-oxazoline L-161,240, the sulfonamide BB-78485 and two methyl sulfones, PF-04753299 and PF-05081090, was analyzed. LpxC inhibitors share common features like a hydroxamic acid moiety (light purple), which chelates the catalytically Zn^2+^ and a large hydrophobic area (light blue dashed), which occupies the hydrophobic binding tunnel of LpxC. By exchanging the hydrophobic residues, a broad range of different chemical classes can block LpxC.

Based on the known amino acid residues required for substrate interaction and enzyme activity, many LpxC inhibitors have been developed, bioinformatically analyzed and intensively tested *in vitro* over the last two decades (22–26). Their antimicrobial activity was screened in various pathogens, e.g. *E. coli*, *Pseudomonas aeruginosa, Neisseria meningitidis* or *Salmonella enterica* (27–29). Structurally, the developed inhibitors share common features such as a hydroxamic acid moiety coordinating the catalytically important Zn^2+^ and a hydrophobic region occupying the lipophilic binding tunnel (30). Among a variety of chemical classes, the *N*-aroyl-L-threonine- and methyl sulfone-based compounds are yet the most promising candidates in terms of LpxC inhibition and antibacterial activity.

Here, we analyzed the effect of five different commercially available LpxC inhibitors on the Gram-negative model bacterium *E. coli*: the *N*-aroyl-L-threonine derivative CHIR-090, the aryl-oxazoline L-161,240, the sulfonamide BB-78485 and two methyl sulfones, PF-04753299 and PF-05081090 (Fig. 1B) (reviewed in Kalinin and Holl, 2016). We compared their far-reaching effects on cell viability, membrane composition and LpxC stability as well as the overall proteomic response to these compounds. This study shows how cells counter the inhibition of LpxC, which will help to understand further regulatory relationships in the tightly controlled membrane homeostasis network.

## 2. Results

### 2.1. All tested LpxC inhibitors bind to purified LpxC

To the best of our knowledge, the five different LpxC inhibitors (Fig. 1B) have never been directly compared, and it is as yet unclear whether they elicit a universal or compound-specific cellular response. Therefore, we started our analysis by showing that they all bind *in vitro* to LpxC under identical assay conditions. Purified His-tagged LpxC was subjected to a thermal shift assay with the cysteine-binding fluorescent probe, BODIPY^TM^ FL L-cystine. When the temperature gradient reaches the melting temperature (T_m_) of the protein, it starts to unfold and exposes its cysteines, which leads to an increased fluorescence signal due to the interaction with BODIPY^TM^ FL L-cystine. After reaching a fluorescence maximum, the protein starts to aggregate, which reduces the signal intensity (Fig. 2A). Binding events like protein-protein or protein-ligand interactions can influence the melting properties of the protein.

**FIGURE 2.**
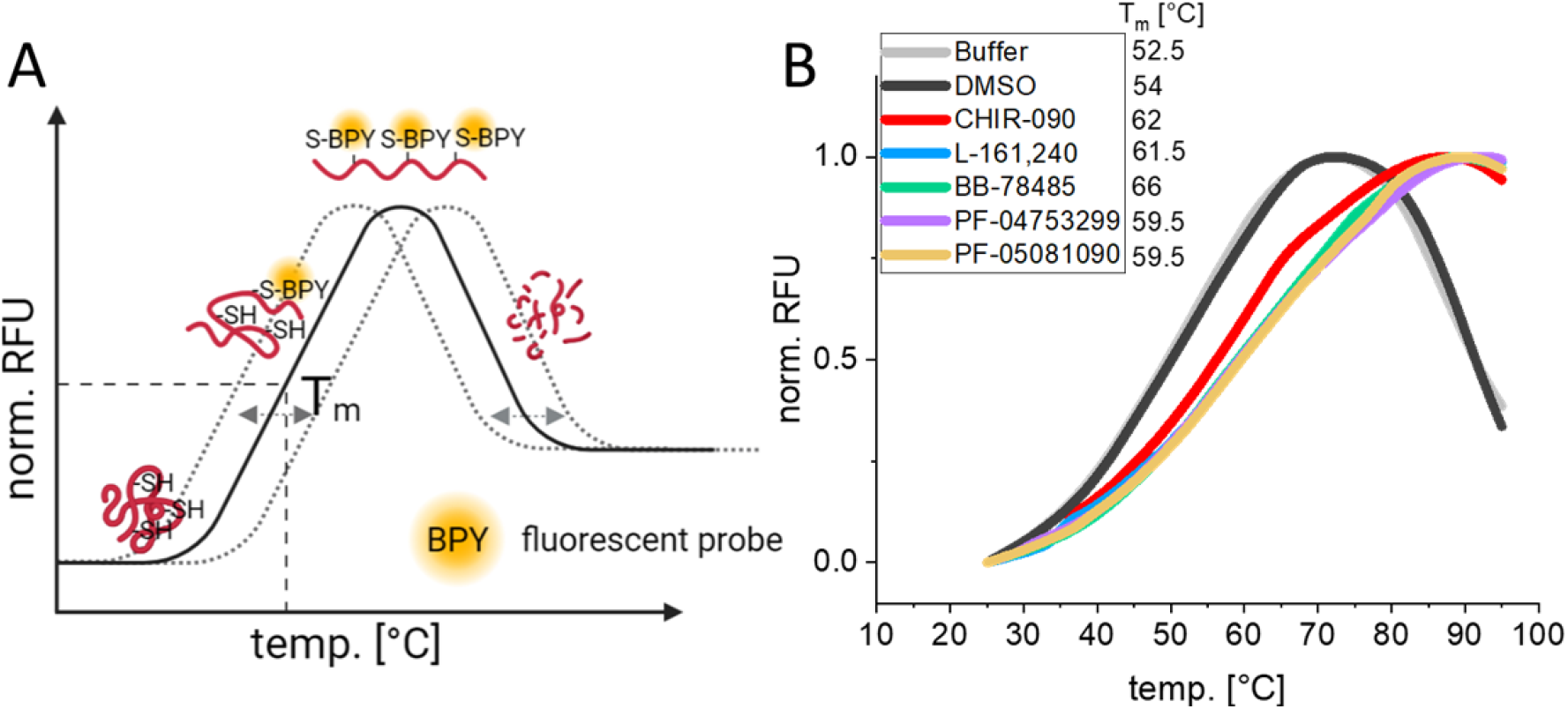
LpxC inhibitors bind to purified LpxC *in vitro*. A) In the thermal shift assay, a fluorescent probe (BODIPY^TM^ FL L-cystine, BPY) is mixed with a protein and subjected to a temperature gradient between 20 and 100°C. With increasing temperature, the protein starts to unfold, and the probe interacts thiol-specifically with the exposed cysteines of the protein. This leads to an increase in fluorescence. At a certain temperature, the protein starts to aggregate, causing the fluorescence signal to decrease. The binding of ligands or protein interaction partners can reduce or increase the melting temperature (T^m^; dotted grey lines). B) 12.5 µM purified His-tagged LpxC with 2 µM BODIPY^TM^ FL L-cystine was assayed without or with 50 µM of the LpxC inhibitors. While the curves of the samples with LpxC plus buffer or the solvent DMSO overlap, all inhibitors caused a substantial shift in the melting point of LpxC. The measurements were performed in technical triplicates and the mean value was used for the graph generation. RFU= relative fluorescence unit.

LpxC in buffer or in buffer with DMSO (in which the LpxC inhibitors were solved) showed a melting temperature of 52.5 or 54°C, respectively (Fig. 2B). Each of the five compounds stabilized the protein by increasing its melting point by at least 7°C, a clear indication that they bound to LpxC.

### 2.2. The membrane is not depolarized by the amphipathic character of LpxC inhibitors

Since all five compounds have a polar moiety and a large hydrophobic region (Fig. 1B), we asked whether they have amphipathic properties that disturb the membrane integrity of living bacteria. It is known that the membrane can be depolarized by amphipathic molecules such as polymyxin B, a cationic polypeptide antibiotic (31). With the help of the fluorescence probe DiSC3, which integrates into polarized, but not depolarized membranes, the loss of membrane potential can be monitored over time (Fig. 3A). In this assay, the fluorescence signal stabilizes after an integration phase. Upon membrane depolarization, e.g., by the addition of polymyxin B, the probe is released, which leads to an increase of fluorescence. While polymyxin B as the positive control induced a rapid upshift in the fluorescence signal, this was not the case after addition of DMSO or either of the five LpxC inhibitors in DMSO (Fig. 3B). This indicates that the structural properties of these compounds have no immediate impact on membrane polarization.

**FIGURE 3.**
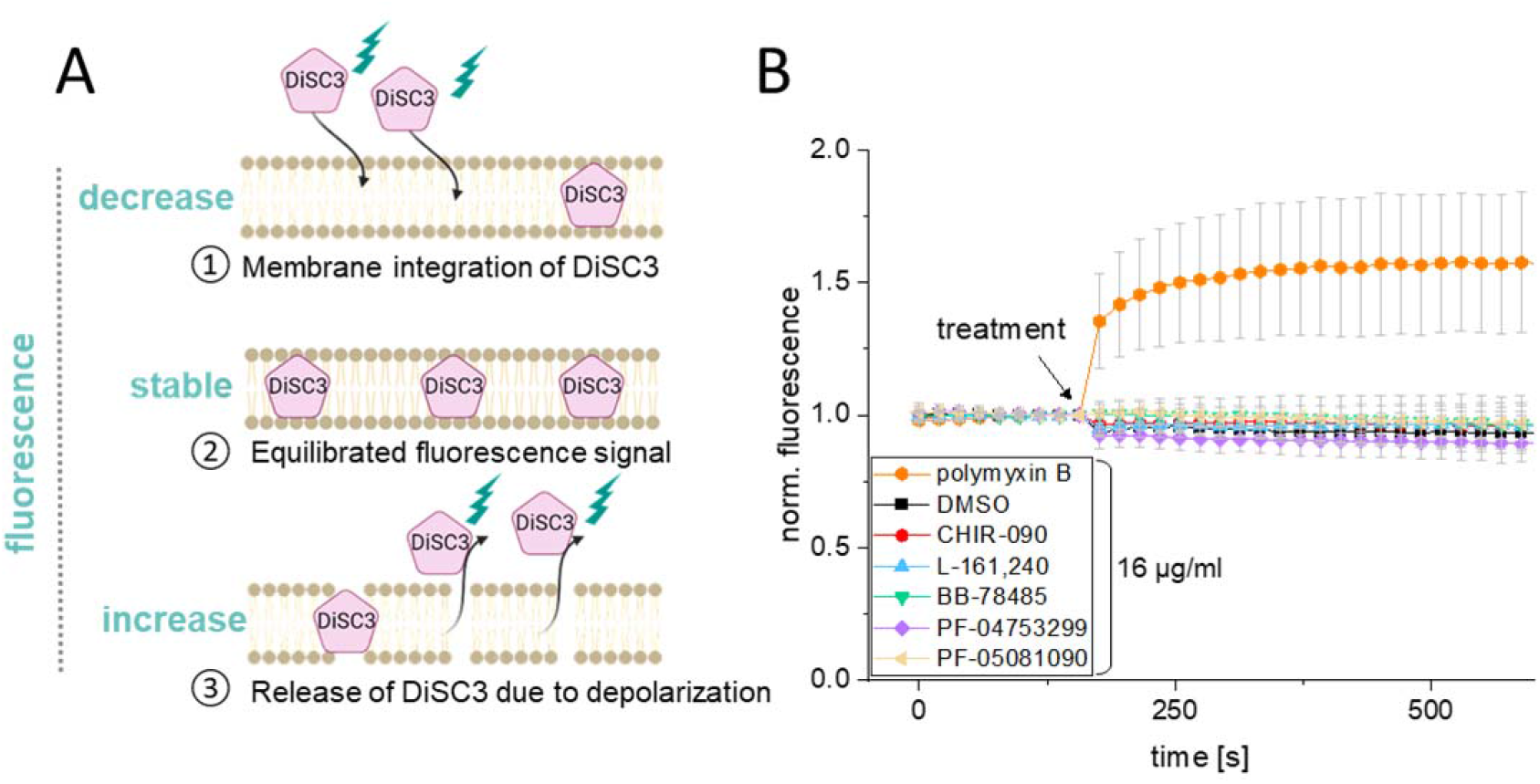
The *E. coli* W3110 membrane is not depolarized by the LpxC inhibitors. Due to their amphipathic structure, it was conceivable that the inhibitors chemically depolarize the bacterial membrane as it is known for polymyxin B. A) DiSC3 is a membrane integrative fluorescence probe, which is used to visualize short-term membrane depolarization as a response to specific treatments. After addition of DiSC3, it starts to integrate into the membrane (①). The fluorescence signal stabilizes upon maximum incorporation (②). Then, the effect of various compounds on membrane integrity can be examined. If the membrane is disturbed, the incorporated DiSC3 is released, thereby leading to an increase of the fluorescence signal (③). B) Exponential *E. coli* W3110 cells were adjusted to an OD_580nm_ of 0.5. After equilibration of the initial fluorescence, cells were exposed to 16 µg/ml of one of the LpxC inhibitors, polymyxin B or, as negative control, to DMSO (final concentration: 1.6 %). The experiment was performed in biological triplicates with each time three technical replicates, standard deviations are indicated and the values before addition of DiSC3 were set to 1 to visualize the change of the fluorescence signal.

### 2.3. Physiological effects in response to treatment with LpxC inhibitors

To be able to compare the bacterial responses to sublethal doses of the five compounds, we determined their relative potency. *E. coli* W3110 growth in M9 minimal medium with different concentrations of the LpxC inhibitors was monitored to obtain the minimal inhibitory concentration (MIC) (Fig. S1). While CHIR-090, L-161,240 and PF-05081090 only required 0.2 µg/ml to inhibit growth, 1 µg/ml of PF-04753299 and 5 µg/ml of BB-78485 were necessary (Table 1). We also determined the MICs in the efflux pump deficient *E. coli* W3110 Δ*tolC* strain (Fig. S1). This strain was eight to ten times more susceptible, suggesting that the wild type can lower the intracellular concentration of LpxC inhibitors by export.

**TABLE 1.**
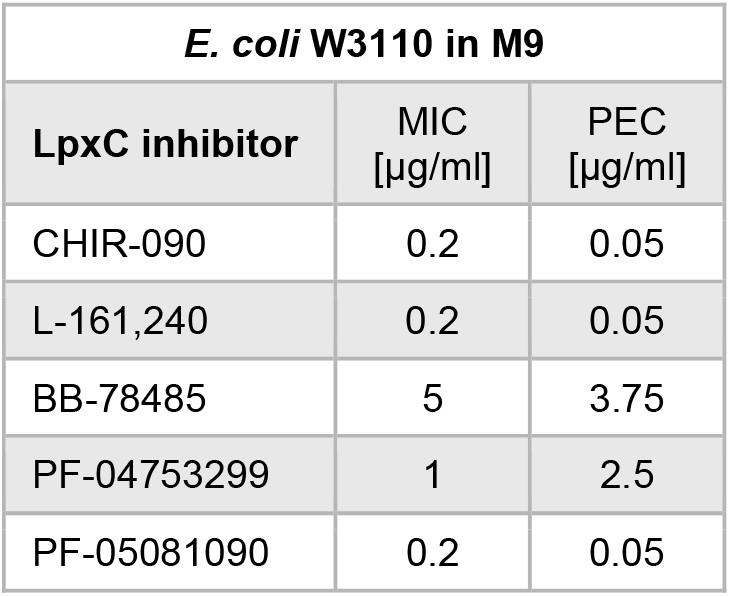
Determination of the minimal inhibitory concentration (MIC) and the physiologially effective concentration (PEC) for the LpxC inhibitors. *E. coli* W3110 cells were grown in M9 minimal medium at 37°C under constant agitation. Compounds were diluted in DMSO (final concentration 0.3 %), which had no effect itself on the growth.

To allow the bacteria to physiologically respond to the compounds, for example in experiments monitoring the proteome response to antibiotic treatment (see below), it was necessary to determine the adequate physiologically effective concentration (PEC). This is defined as the concentration required to inhibit growth by at least 30 % within the first two hours after addition of the compound. Only ¼ of the MIC was required for four of the inhibitors (Fig. S2A). Higher concentrations resulted in a decrease in the optical density, suggesting cell lysis. Compound BB-78785 appeared to be less potent, and ¾ of the MIC was defined as the PEC (Table 1).

As the LpxC inhibitors are supposed to block an essential pathway, we examined the cell morphology and viability after treatment using microscopy. On average, *E. coli* cells became shorter and wider in the presence of 1.5 x MIC after five hours. The majority of the treated cells was about 1 µm shorter, and this was accompanied by a slight increase in width giving the bacteria an overall rounder appearance (Fig. 4A,B). Staining with the DNA-binding dye propidium iodide (PI) provided evidence for a disturbed membrane integrity in these visibly altered bacteria (Fig. 4C). PI only penetrates the cell when the membrane is partially or completely disrupted. While hardly any PI-stained cells were found five hours after DMSO addition, many cells were strongly fluorescent after inhibitor-treatment indicating that the dye had access to the cytoplasm. Calculating the relative number of unaffected bacteria on the basis of these microscopy results showed that more than 98% of the cells remained intact in the presence of DMSO. In contrast, all five LpxC inhibitors had a severe effect, ranging from about 50% cells remaining intact after BB-78485 or PF-04753299 treatment to less than 30% in the case of CHIR-090, L-161,240, or PF-05081090 (Fig. 4C).

**FIGURE 4.**
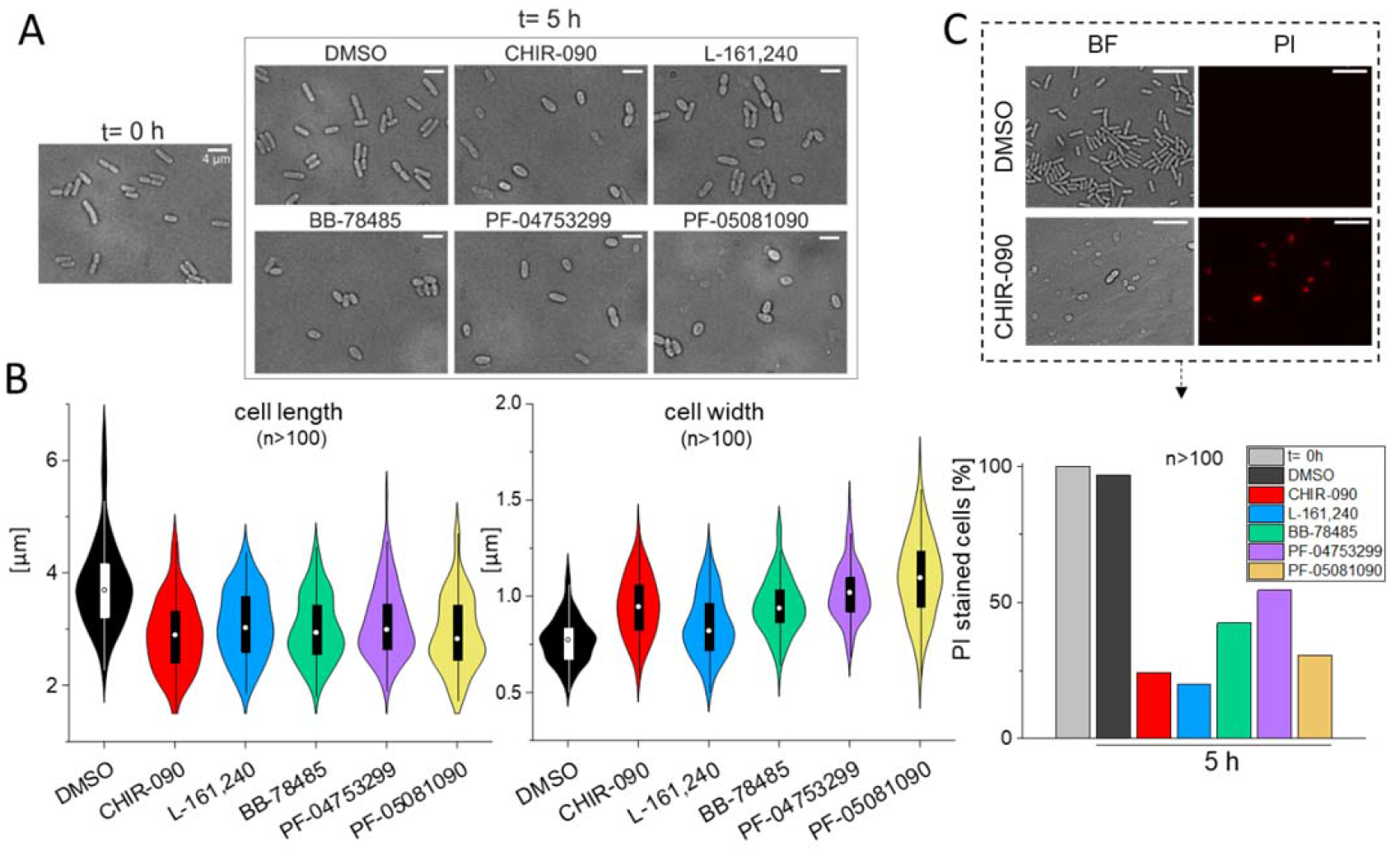
Cell shape and viability are affected by LpxC inhibitors. A) Brightfield microscopy of cells cultivated in M9 medium after five hours of exposure to LpxC inhibitors (1.5x MIC). DMSO-treated control cells retained their rod shape, whereas cells were shorter and wider in the presence of LpxC inhibitors. B) Cell length and width quantification (n>100) with the software ImageJ. C) Changes in the cell shape correlate with higher uptake of the propidium iodide (PI) stain. Bacteria were stained with (PI) after 5 hours of exposure to 1.5x MIC of the inhibitors. This dye only enters the cell when membrane integrity is impaired. As representative example, the ratio of total cells counted to cells stained with PI was determined (n>100), and the membrane integrity of the cells before addition of the compounds was set to 100%. Scale bar (A)= 4 µm, Scale bar (C)= 8 µm

These results suggested a defect in membrane homeostasis triggered by the LpxC inhibitors. To follow up on this assumption, we examined the susceptibility to these compounds in combination with other stressors targeting the cell envelope. As a control we used the antibiotic spectinomycin, which blocks translation by binding to the 30S ribosome. After two hours of incubation with DMSO, the LpxC inhibitors or spectinomycin, cells were washed, adjusted to the same optical densities, serially diluted, and spotted onto agar plates. No growth defect was observed on LB plates upon pre-treatment with the LpxC inhibitors alone (0.25 x MIC; Fig. 5, two further replicates in Fig. S3). Apparently, severe growth defects were prevented by removing the compounds by washing followed by cultivation on rich medium. The presence of ampicillin or SDS + EDTA in the plate had no or little effect, respectively. The most severe effect after prior exposure to the LpxC inhibitors was observed for vancomycin. This glycopeptide antibiotic targeting cell wall biosynthesis can only cross the OM of Gram-negative bacteria when the latter is compromised.

**FIGURE 5.**
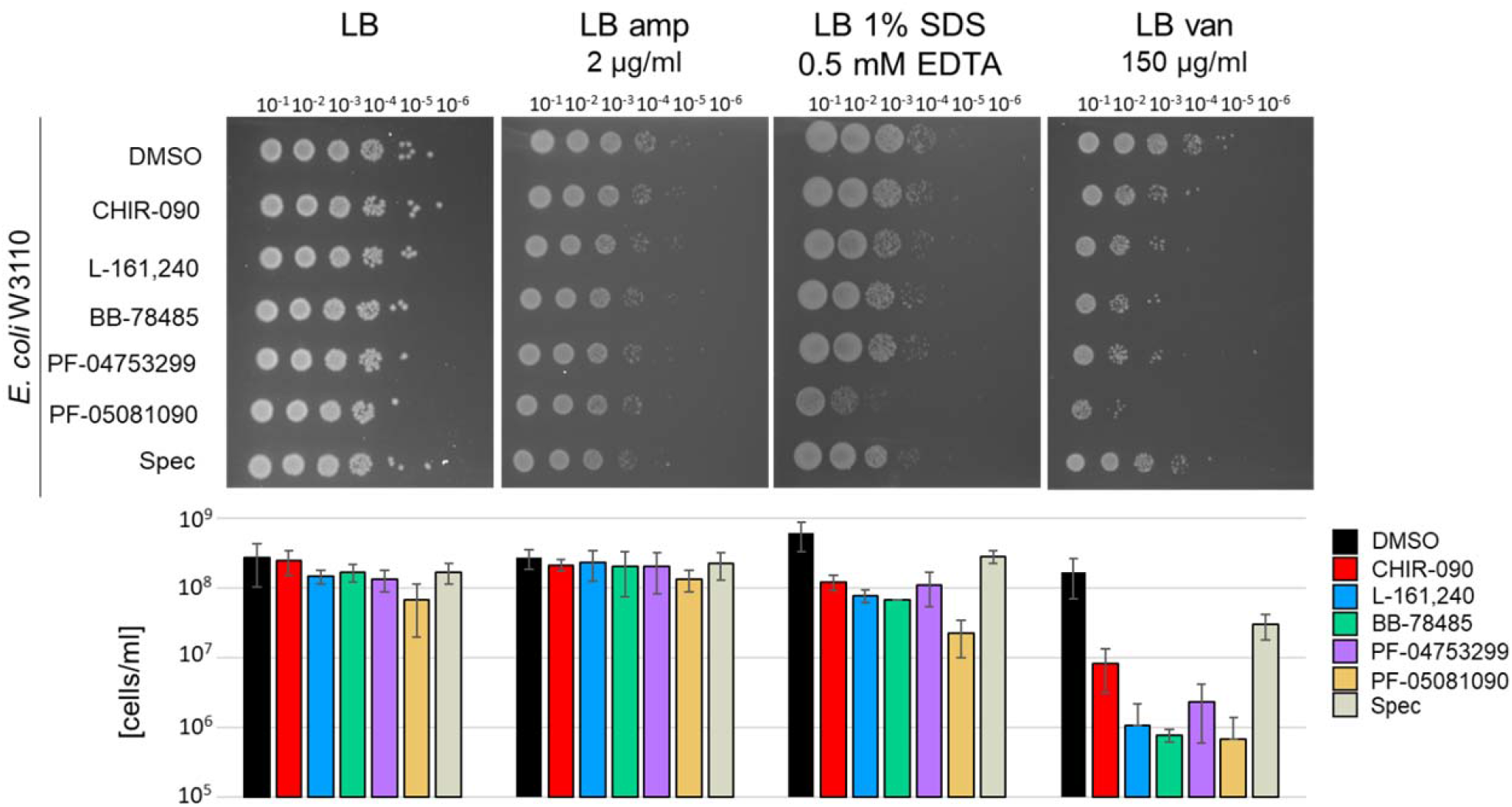
Susceptibility of *E. coli* W3110 pre-treated with LpxC inhibitors to stressors targeting the cell envelope. Sensitivity toward cell envelope-active antibiotics such as ampicillin (amp) or vancomycin (van) and the membrane-disrupting detergent SDS was analyzed. Washed pre-treated *E. coli* W3110 cells (exposed to ¼ MIC of the inhibitors or, as control, to 300 µg/ml spectinomycin (spec) for two hours in LB medium) were diluted from 10^-1^ to 10^-6^ in 0.9% NaCl and then spotted onto different agar plates (LB, LB amp, LB SDS/EDTA and LB van). After overnight incubation at 37°C, pictures were taken (upper panel). The number of surviving cells per ml from three biological triplicates (Fig. S3) was calculated by counting the colonies of the spot with highest dilution (lower panel).

### 2.4. Effects of LpxC inhibitors on LpxC levels and membrane composition

Next, we wondered whether the cellular LpxC and LPS levels changed in response to LpxC inhibitors. Since the five inhibitors elicited largely similar effects, we used CHIR-090 as model compound for this experiment. Cell growth was followed for five hours, and a sample was taken every hour and prepared for SDS-PAGE analysis to determine the LpxC and LPS levels by Western blot analysis. Compared to cells exposed to DMSO alone, immediate growth suppression was observed after addition of spectinomycin (Spec), while cells treated with CHIR-090 grew for almost two hours before they started to lyse (Fig. 6A). Treatment with CHIR-090 but not DMSO or Spec induced a strong increase in LpxC abundance (Fig. 6B, triplicates in Fig. S4). Elevated LpxC levels were not accompanied by elevated LPS levels suggesting that binding of the inhibitor to LpxC (Fig. 2) inactivates the enzyme.

**FIGURE 6.**
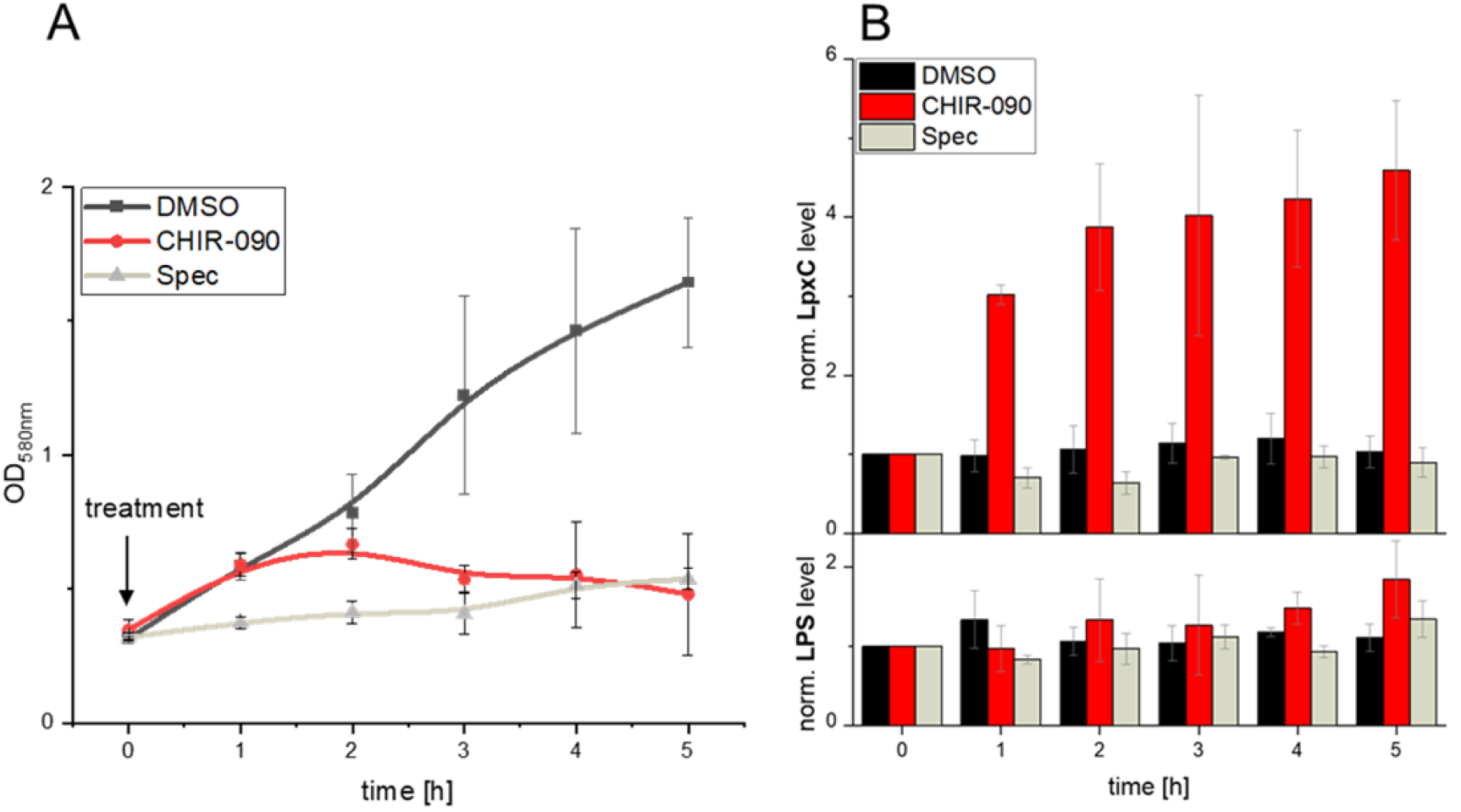
LpxC levels increase in response to CHIR-090, whereas the LPS levels remains relatively constant. A) The growth of exponential *E. coli* W3110 in M9 minimal medium after addition of DMSO, CHIR-090 (2 x MIC; 0.4 µg/ml) or spectinomycin (spec; 300 µg/ml) was tracked for five hours. Growth inhibition was observed for cells exposed to CHIR-090 and Spec. B) Samples were taken every hour and directly frozen in liquid nitrogen. After harvesting, the cell pellets were resuspended to the same optical density in TE-buffer and loading dye. 15 µl of each sample was subjected to SDS-PAGE followed by western transfer on nitrocellulose. Via fluorescent immunodetection, LpxC and LPS levels were quantified with the software ImageLab (BioRad). The experiment was performed in triplicates (Fig. S4).

Various transcriptional or posttranscriptional mechanisms may account for the induction of LpxC upon CHIR-090 treatment. Since CHIR-090 does not affect *lpxC* transcription (32), the most likely explanation is that binding of the compound prevents degradation of LpxC by the FtsH protease. To test this hypothesis, we measured the half-life of LpxC in stationary *E. coli* W3110 cells treated with CHIR-090 or other LpxC inhibitors. After inhibiting protein biosynthesis by adding chloramphenicol, LpxC levels were monitored over time by Western blot analysis with LpxC-specific antisera.

As expected, the LpxC signal decreased in DMSO-supplemented control cultures (Fig. 7A). The addition of all five LpxC inhibitors resulted in a strong increase of LpxC within the 15-minute period between the addition of the inhibitor and chloramphenicol. Furthermore, the signal remained stable for at least two hours, which clearly indicates a protection of LpxC from FtsH-mediated proteolysis (Fig. 7B). To analyze whether inhibition of LPS biosynthesis had an impact on the connected PL biosynthesis (Fig. 1A), we extracted the total phospholipid species from inhibitor-treated bacteria and subjected them to thin layer chromatography (TLC). Molybdenum blue staining visualized the most prominent PL species of *E. coli,* namely phosphoethanolamine (PE), phosphatidylglycerol (PG) and cardiolipin (CL). Except for BB-78485, a new spot appeared after treatment with the other four compounds (Fig. 8A, two replicates in Fig. S5A). By comparison with lipid standards, the new PL most likely is lyso-PE (LPE). Ninhydrin, which stains free amino groups available in lipids like phosphatidylserine (PS), PE and LPE, detected the same spots fully supporting this hypothesis. Moreover, LPE has previously been seen after treatment of *E. coli* with a larger excess (4x MIC) of CHIR-090 (33). LPE can result from PE cleavage by two alternative pathways: either by the OM-anchored phospholipase PldA, which serves as a sensor for lipid asymmetry, or by PagP, which also resides in the OM and transfers a palmitate chain from a PL to produce a hepta-acylated lipid A molecule (34, 35). To distinguish between these two possibilities, we used the Δ*pldA* and Δ*pagP* mutants from the Keio collection (36). Exposure to CHIR-090 triggered LPE production in the Keio WT strain and the Δ*pagP* mutant, but not in the Δ*pldA* mutant (Fig. 8B, two further replicates Fig. S5B). We conclude that LPE accumulation in response to four of the five LpxC inhibitors is attributable to PldA suggesting a misbalance in membrane homeostasis.

**FIGURE 7.**
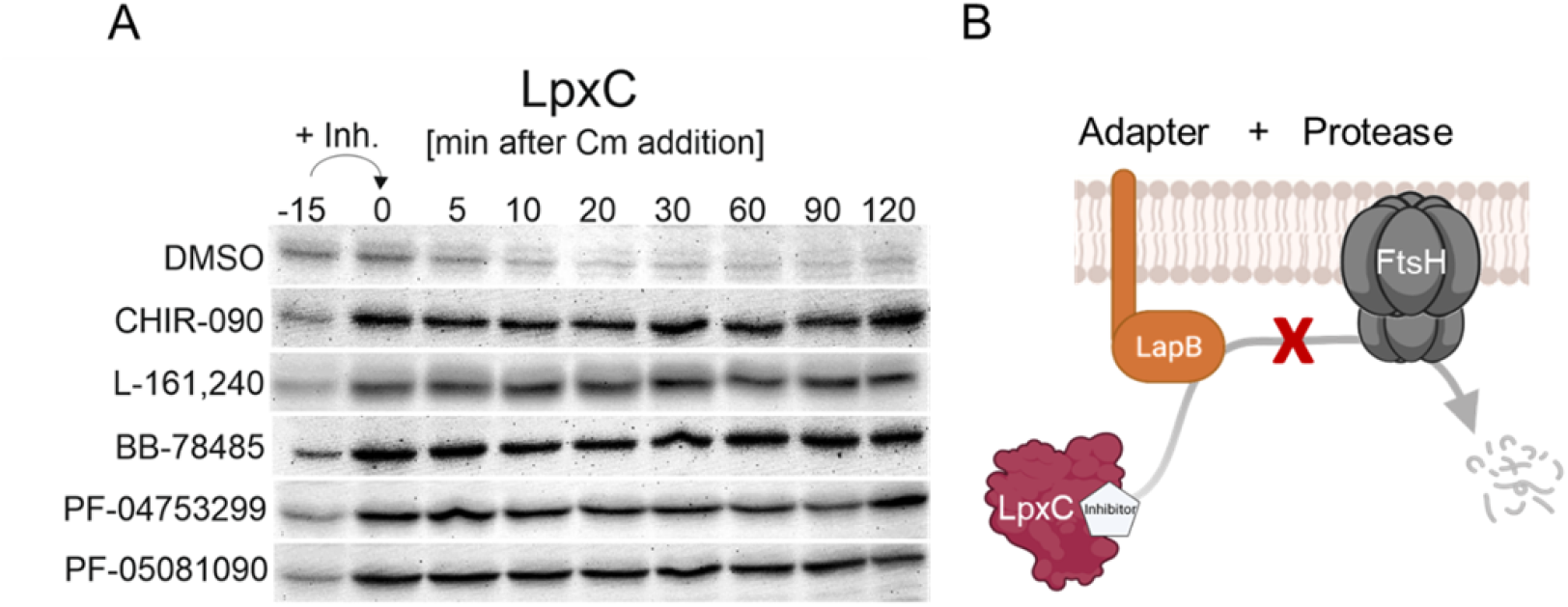
LpxC stability increases upon addition of LpxC inhibitors. A) Chromosomally encoded LpxC levels were determined by in vivo degradation experiments. Stationary *E. coli* W3110 cells were exposed to LpxC inhibitors for 15 min before 200 µg/ml chloramphenicol (Cm) was added to inhibit protein biosynthesis. The determined MICs of each inhibitor were used. 1 ml samples were drawn before addition of the inhibitor (−15), before addition of Cm (0) and after 5, 10, 20, 30, 60, 90 and 120 min. The protein contents were adjusted by resuspension of the cell pellets according to their optical density (100 µl for OD_580_= 1) in TE buffer and loading dye. 15 µl of each sample were separated by SDS-PAGE and blotted onto a nitrocellulose membrane. LpxC was detected by chemiluminescence after incubation with a polyclonal LpxC antiserum and an HRP-coupled goat anti-rabbit antibody. The experiment was performed in triplicates with similar results. B) The LapB/FtsH-mediated turnover of LpxC appears to be reduced in the presence of LpxC inhibitors. This could either be a physiological response (consequence of blocking LPS synthesis) or indicate impaired interaction with LapB or prevented FtsH recognition.

**FIGURE 8.**
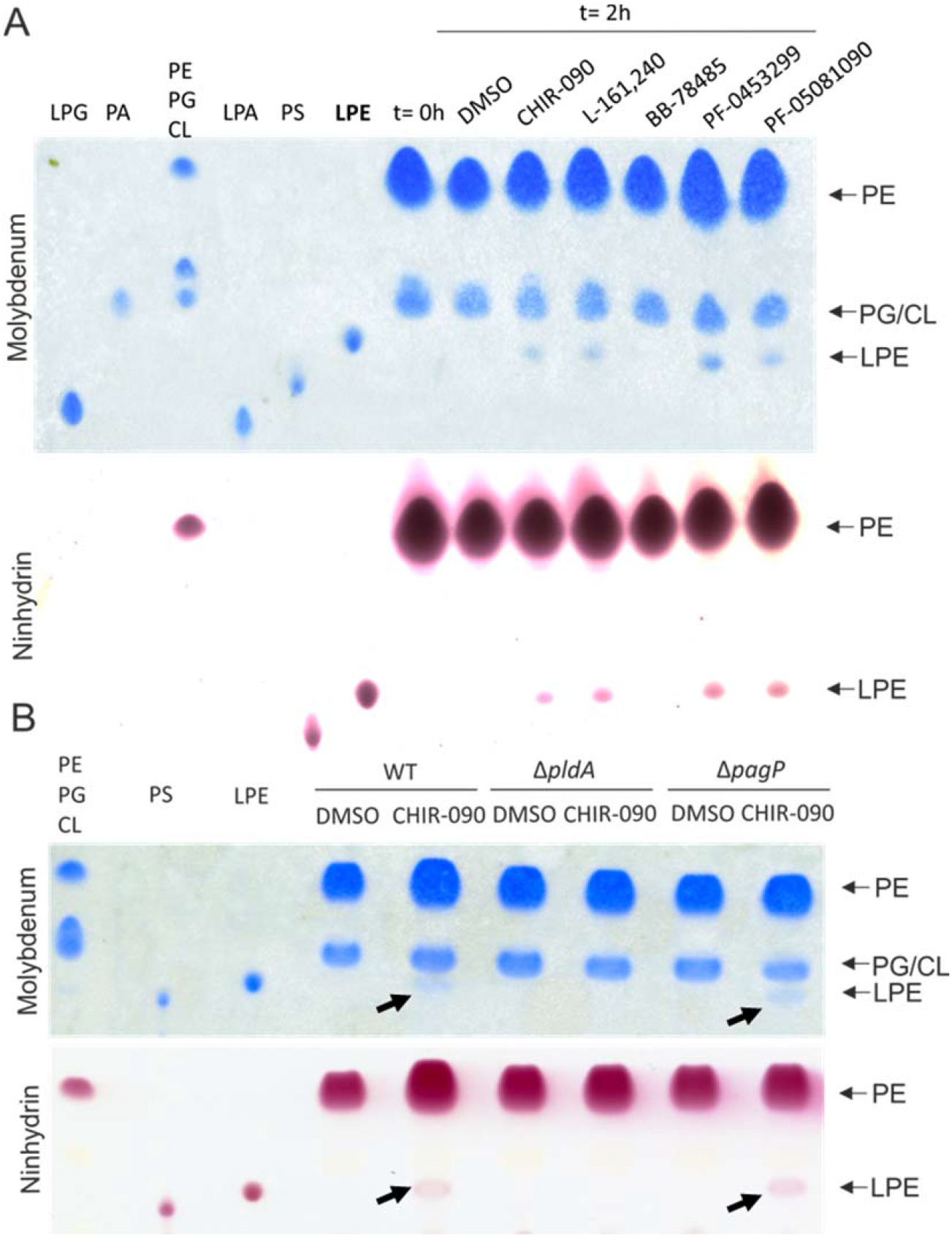
Four of the five LpxC inhibitors induce PldA-mediated LPE accumulation. A) *E. coli* W3110 was cultivated in M9 minimal medium till exponential phase before the culture was divided (t= 0h) and treated with DMSO or ½ MIC of the LpxC inhibitors for 2 h. Lipids from cell pellets according to OD_580nm_=10 in 1 ml were extracted and separated via TLC using chloroform:methanol:water (65:25:4) as mobile phase. Phospholipids were visualized by molybdenum blue spray reagent and lipids with free amino groups were stained with a ninhydrin staining solution. The retention behavior of commercially available phospholipids is shown to the left. B) *E. coli* BW25113 (WT) and the corresponding Keio-mutants Δ*pldA* and Δ*pagP* were cultivated in M9 minimal medium till exponential phase, then the culture was divided (t= 0h) and treated with DMSO or ½ MIC of CHIR-090 for 2 h. Lipid extraction and TLC analysis was performed as described in A). The experiments were performed in biological triplicates (Fig. S5). LPG: lyso-phosphatidylglycerol; PA: phosphatidic acid; PE: phosphatidylethanolamine; PG: phosphatidylglycerol; CL: cardiolipin; LPA: lyso-phosphatidic acid; PS: phosphatidylserine; LPE: lyso-phosphatidylethanolamine.

### 2.5. The proteomic response to LpxC inhibitors

Hitherto, the obtained results indicate a multi-faceted response of *E. coli* to LpxC inhibition. To gain insights into the commonalities and differences between the responses to the individual compounds, we monitored the global changes at the proteome level by using radioactive pulse-labeling and 2D gel electrophoresis. *E. coli* cultures were treated with the previously determined PECs of LpxC inhibitors (refer to table 1), and growth was followed for three hours to ascertain the effectiveness of the compounds (Fig. S2B). Incorporation of L-[^35^S]-methionine into newly synthesized proteins was started 10 min after inhibitor addition for 5 min. Upregulated proteins (at least 1.6 x fold upregulated in each biological triplicate) were identified by comparing relative signal intensities for proteins detected on autoradiographs of 2D gels of inhibitor-treated versus DMSO-treated autoradiographs.

The number of regulated proteins depended on the inhibitor used and varied from eight to over 40 (Fig. 9A-E). Most obviously, BB-78485, which was applied in a 15- to 75-fold higher concentration than the other inhibitors due to its much higher MIC and PEC (Table 1), induced the broadest response (Fig. 9C). It also affected protein synthesis rates to a greater extent than the other compounds (Fig. 9F) although growth of the cultures used for protein labeling was not severely compromised (Fig. S2B).

**FIGURE 9.**
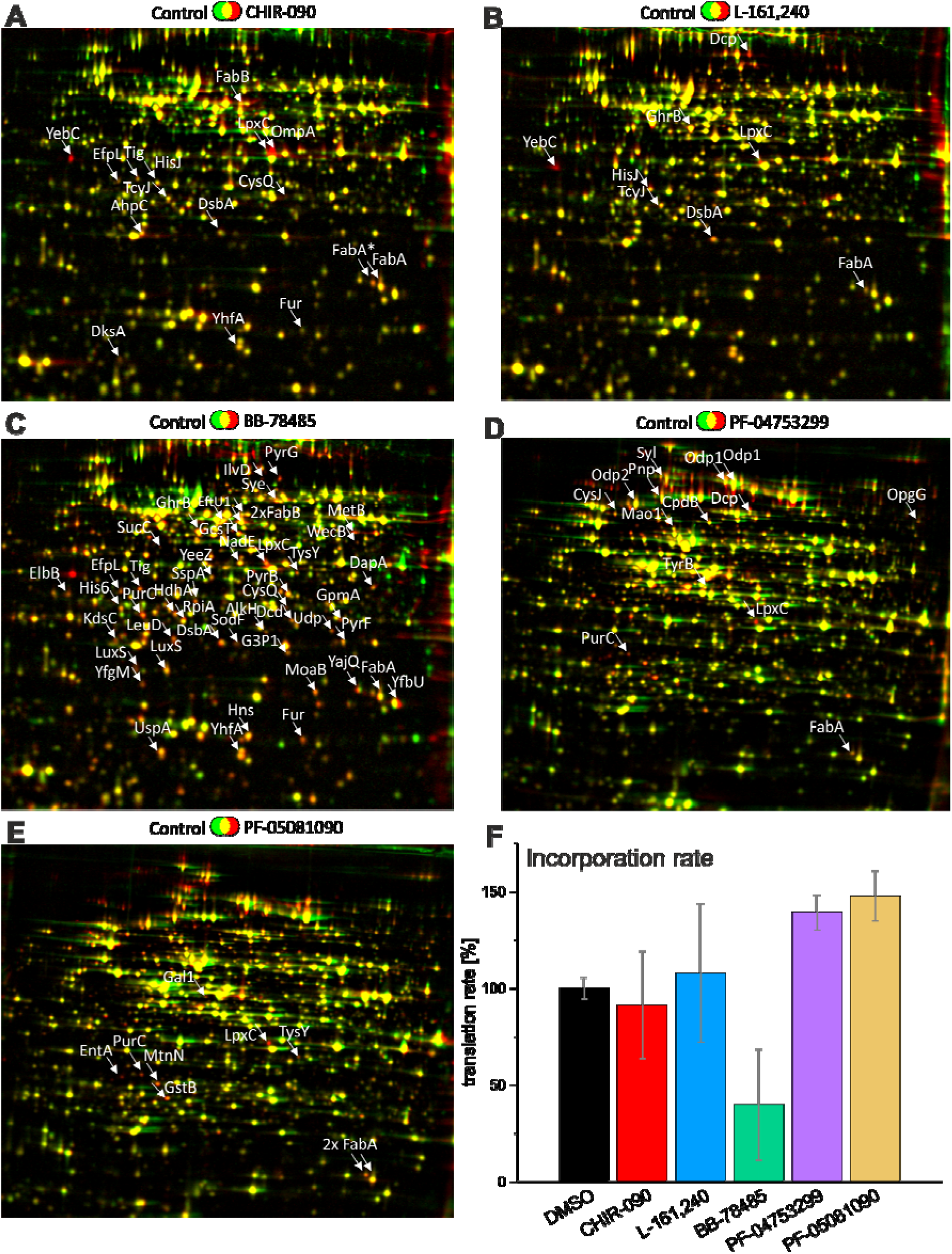
The proteomic response to LpxC inhibitor exposure varies but consistently shows upregulation of LpxC and enzymes of unsaturated fatty acid biosynthesis. *E. coli* W3110 grown in M9 minimal medium until midlog phase was treated with the five LpxC inhibitors at the previously determined PECs. After 10 min, L-[^35^S]-methionine was added to label newly synthesized proteins radioactively for 5 min. After separating the proteins first by isoelectric point and then by molecular weight, the 2D gels were dried, and the labeled proteins were detected using autoradiography. The experiment was performed in biological triplicates. Protein spot intensities were quantified in Delta2D (Decodon). Proteins indicated by arrows are at least 1.6 x fold upregulated in comparison to the control in each triplicate. Upregulated proteins in response to CHIR-090 (A), L-161,240 (B), BB-78485 (C), PF-04753299 (D), and PF-05081090 (E) are indicated. (F) To analyze the effect of the inhibitors on the overall protein biosynthesis, the incorporation rates of five min radioactive L-[^35^S]-methionine pulse-labeling was measured with a scintillation counter. The DMSO control was set to 100% and the values were calculated from technical duplicates of three biological replicates.

Identification by mass spectrometry of the altered marker proteins for each compound (Fig. 9) revealed partially overlapping and partially distinct response patterns (Table 2, Table S1). In agreement with an increase in LpxC levels by stabilization of the protein (Fig. 6B and 7A), LpxC was consistently found upregulated in the 2D gels with some statistical variation (Table 2, Table S1). In agreement with a crosstalk between the LPS and PL pathways (18), at least one of the key enzymes of the fatty acid (FA) biosynthesis, FabA or FabB, increased after exposure to each of the LpxC inhibitors. Other upregulated proteins were specific to one or several inhibitors and fell into different functional categories. Treatment with some of the compounds induced proteins coping with membrane-related or redox stress, like DsbA or YfgM. LuxS, an enzyme required for the synthesis of autoinducer 2 (AI-2), is part of the quorum sensing pathway, and was found to be regulated in several cases. Nucleotide biosynthesis may also be affected, as PurC and PyrF, enzymes involved in purine and pyrimidine synthesis, respectively, were found to be regulated. HisJ or TcyJ and other proteins involved in amino acid uptake and protein metabolism also increased in response to more than one of the compounds suggesting some far-reaching effects into cellular metabolism.

**TABLE 2.** Overview of the proteome responses to LpxC inhibitors. LpxC accumulated in response to all inhibitors. FabA and/or FabB synthesis were upregulated in most samples. Colored fields indicate that the respective protein was upregulated at least 1.6-fold in all replicates (for details, see Table S1). FA: fatty acid, n.d.: not determined.

| proteins | CHIR-090 | L-161,240 | BB-78485 | PF-04753299 | PF-05081090 | protein function |
| --- | --- | --- | --- | --- | --- | --- |
| **membrane synthesis** |  |  |  |  |  |  |
| FabA | 3 ± 0.6 | 2.7 ± 0.7 | 2.9 ± 0.6 | 2 ± 0.1 | 2.2 ± 0.7 | FA synthesis |
| FabB | 2.2 ± 0.3 | 1.6 ± 0.4 | 2 ± 0.2 | 2.9 ± 1.2 | 2.9 ± 1.5 | FA synthesis |
| LpxC | 2.8 ± 0.8 | 1.8 ± 0.2 | 4.7 ± 0.5 | 3.5 ± 1.2 | 2.8 ± 1.8 | LPS synthesis |
| OmpA | 1.8 ± 0.2 | 1.2 ± 0.3 | 2 ± 0.6 | n.d. | n.d. | membrane integrity |
| **stress response** |  |  |  |  |  |  |
| DsbA | 2.1 ± 0.5 | 2.4 ± 0.4 | 2.6 ± 0.7 | 1.5 ± 0.5 | 1.4 ± 0.4 | redox response |
| LuxS | 1.8 ± 0.5 | 1.8 ± 0.5 | 3.3 ± 1.0 | n.d. | n.d. | quorum sensing |
| YfgM | 1.6 ± 0.4 | 2 ± 1.0 | 2.6 ± 0.4 | n.d. | n.d. | chaperone |
| **DNA and energy metabolism** |  |  |  |  |  |  |
| PurC | 1.5 ± 0.2 | 1.3 ± 0.2 | 3.4 ± 0.7 | 2.6 ± 0.8 | 2.6 ± 0.9 | purine biosynthesis |
| PyrF | 1.7 ± 0.1 | 1.4 ± 0.4 | 2.1 ± 0.1 | n.d. | n.d. | pyrimidine biosynthesis |
| **protein synthesis** |  |  |  |  |  |  |
| HisJ | 2.4 ± 0.7 | 2.4 ± 0.2 | 1.3 ± 0.3 | 1.8 ± 0.9 | 1.6 ± 0.7 | amino-acid transport |
| Syl | n.d. | n.d. | n.d. | 1.9 ± 0.4 | 3.1 ± 2.2 | protein biosynthesis |
| TcyJ | 2.4 ± 0.9 | 2.6 ± 0.4 | 1.8 ± 0.6 | n.d. | n.d. | amino-acid transport |
| YebC | 3.1 ± 1.8 | 7.4 ± 7.5 | 6.7 ± 7.0 | n.d. | n.d. | transcription regulation |
| **Other** |  |  |  |  |  |  |
| YhfA | 2 ± 0.3 | 1.8 ± 0.5 | 3.6 ± 1.1 | n.d. | n.d. | unknown function |

## 3. Discussion

LPS biosynthesis is a Gram-negative specific cellular process that emerges as a prime antibiotic target. The major focus lies on LpxC, the broadly conserved committed enzyme essential for lipid A biosynthesis (37). Hundreds of structurally diverse LpxC inhibitors have been synthesized by academic and industrial researchers in the last 20 years (38), and these efforts continue until now (39, 40). *In silico* docking studies, *in vitro* binding assays and *in vivo* MICs for clinically relevant Gram-negative bacteria have been reported for many of these compounds, many of which show promising MIC values in the nanomolar or low micromolar range(41, 42). So far, only a few LpxC inhibitors entered first clinical trials where they failed due to inflammation at the injection sites or dose-limiting cardiovascular toxicity (30, 43).

### 3.1. Similar yet different responses of *E. coli* to LpxC inhibition

Despite intense efforts directed towards the discovery of novel LpxC inhibitors, surprisingly little is known about the cellular responses of Gram-negative bacteria to such compounds. Here, we compared the physiological and proteomic response of *E. coli* to five different LpxC inhibitors, which elicited similar but not identical responses. The chemical properties of neither of these structurally heterogeneous compounds (Fig. 1B) had an immediate impact on membrane integrity as shown by a membrane depolarization assay (Fig. 3) suggesting that they reach the cytoplasm and exert their activity from the inside. This is supported by an 8- to 10-fold lower MIC in the efflux pump-deficient strain *E. coli* W3110 Δ*tolC*. Like many other antimicrobial compounds (44), the LpxC inhibitors seem to be exported by multi-drug efflux pumps. This was also shown to be case for CHIR-090 in *P. aeruginosa* (45, 46).

Over the course of several hours, all inhibitors induced cell shrinkage, higher susceptibility to vancomycin and increased uptake of propidium iodide, providing evidence that the membrane integrity was compromised. LpxC levels went up and stayed stable upon protein synthesis inhibition indicating that inhibitor-bound enzyme is (i) inactive because LPS levels remained low and (ii) not degraded by the FtsH protease. The latter can probably be explained by the inaccessibility of recognition motifs in LpxC (14, 47), the inability to interact with the machinery that delivers LpxC to FtsH (48–50), or changes in metabolites, e.g. lipid A disaccharide, acyl-CoA or periplasmic LPS intermediates, affecting LpxC stability (19, 51, 52).

Instead of LPS, the phospholipid LPE was induced by four of the five inhibitors. It was generated by PldA, a phospholipase in the OM with an established role in signaling perturbations in membrane homeostasis(51). PldA catalyzes the hydrolysis of acyl ester bonds in phospholipids at the outer leaflet of the OM to remove mislocalized PL. The alternative hypothesis, that the lipid A palmitoyltransferase PagP was the LPE producer, was not supported because LPE formation occurred in the *pagP* mutant. This is consistent with the previous finding that deletion of *pldA* reduces the viability of a LPS misregulated mutant, while deletion of *pagP* had no effect (53). In summary, the first adaptation mechanism found in response to four of the LpxC inhibitors was the maintenance of PL homeostasis by the action of PldA.

The second common adaptation mechanism was revealed by 2D gel-based proteomics. It also is linked to membrane homeostasis and concerns the biosynthesis of unsaturated fatty acids (UFAs). After exposure to the compounds, FabA, the key enzyme of the UFA biosynthesis, was upregulated, often together with FabB, another enzyme of the fatty acid elongation cycle (Fig. 10). As LpxC inhibition leads to an accumulation of saturated acyl-ACP, FabA overproduction is probably required to maintain a balanced pool of UFA and SFA. The transcription of *fabA* and *fabB* was previously shown to be stimulated after exposure to CHIR-090 in a FabR (fatty acid biosynthesis regulator) dependent manner that senses the UFA:SFA ratio (32). Such an adaptation of lipid composition could alter membrane fluidity and was observed also under other membrane stress conditions like the presence of octanoic acid or solvents, like isobutanol (54, 55). Another regulatory link between the LPS and PL synthesis is the activation of LpxK by UFA (5, 56). This finding, together with the observed induction of KdsC, an enzyme of the Kdo synthesis pathway, suggests an impact on LPS maturation by LpxC inhibitors (Fig. 10).

**FIGURE 10.**
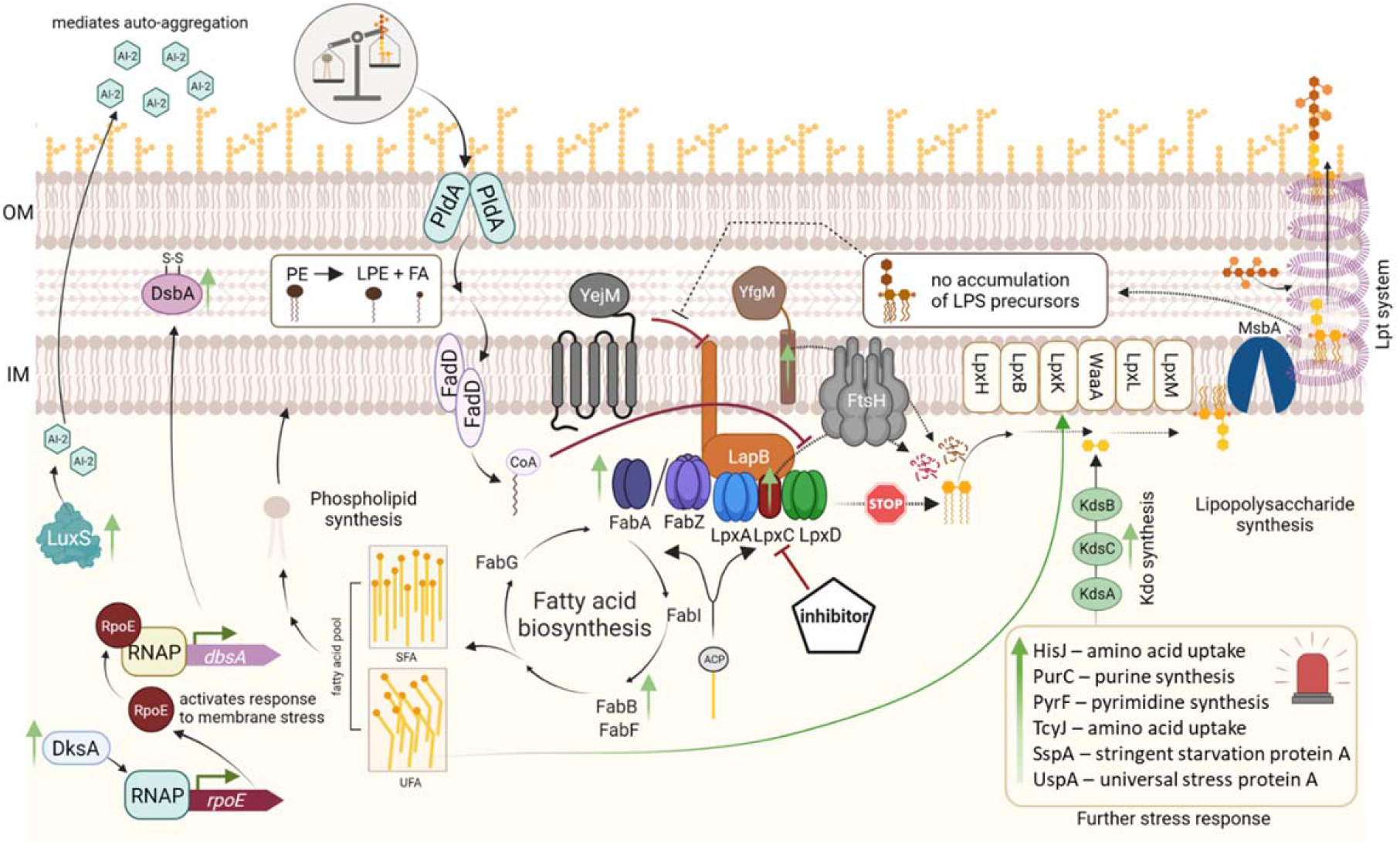
General response of *E. coli* to the inhibition of LpxC, the key enzyme of LPS biosynthesis. When LpxC is inhibited, LPS biosynthesis is blocked (STOP). In turn, lipid A-containing LPS precursors do not accumulate in the periplasm, which stabilizes the YejM/LapB interaction and thereby reduces LpxC turnover by FtsH. The cell likely detects an imbalance of PL and LPS in the outer membrane through PldA, initiating translocation of PE from the outer membrane. Cleavage of PE leads to the accumulation of LPE and fatty acids (FA), which are converted to acyl-CoA by FadD at the cytoplasmic site of the inner membrane. Acyl-CoA has also been shown to stabilize LpxC (51). As a defense mechanism, several proteins are upregulated (indicated by green arrows pointing upwards). These proteins are involved in either stress signaling or membrane synthesis. FabA, key enzyme of the unsaturated FA synthesis, together with FabB, could convert the accumulating acyl-ACP precursors, leading to an increase in unsaturated FA. This would ensure the maintenance of the optimal UFA:SFA (saturated FA) ratio. UFA were also shown to activate LpxH (5) and along with increased Kdo synthesis, this could promote LPS maturation. The increased formation of LuxS is likely to produce autoinducer 2 (AI-2). This is a signal molecule, important for cell-cell communication via quorum sensing, and could mediate growth reduction and auto aggregation within the population. Increased abundance of the global regulator DksA induces *rpoE* transcription. This sigma factor is induced under membrane stress conditions and stimulates the expression *dsbA* leading to increased levels of DsbA. YfgM is an FtsH substrate involved in membrane stress response.

Apart from membrane biogenesis-related responses, LpxC inhibitors elicited several metabolic adaptations (Fig. 10). Similar to the stringent response, LpxC inhibition led to an upregulation of several proteins involved in amino acid synthesis or uptake (Table S1) (57). Treated bacteria probably tried to combat growth limitation by an enhanced uptake of amino acids by proteins like HisJ or TcyJ. Another hint for a global metabolic shutdown was the CHIR-090-induced increase of DksA, which was also regulated in *P. aeruginosa* in response to CHIR-090 exposure (45). DksA is a transcriptional regulator that synergistically binds together with the alarmone ppGpp to RNA polymerase, thereby modulating gene expression in response to amino acid starvation (58). Upon exposure to BB-78485, the nutrient-deficiency regulators SspA (stringent starvation protein A) and UspA (universal stress protein A) accumulated, which also suggest a “starvation-like” feedback (59, 60).

Induction of LuxS by several inhibitors might suggest a population-wide stress signaling by quorum sensing (QS), which can induce auto-aggregation and biofilm formation (61). Aggregation in response to the inhibitors was indeed observed under the microscope (Fig. S7). This can prevent antibiotics but also nutrients from entering single cells (62) and is consistent with the finding that the treated bacteria shut down their metabolism. LuxS-produced autoinducer-2 (AI-2) accumulates primarily during the shift from exponential to stationary phase (63). Inhibition of QS was shown to lower the virulence of bacterial infections (64). Combinatorial approaches that quench OS while inhibiting LpxC might prevent cells from communicating and adapting their metabolism.

Oxidative stress defense proteins, like AhpC, CpdB, SodF and DsbA, were also activated by several LpxC inhibitors. Some of them might be induced through the extracytoplasmic stress response by the sigma factor RpoE, and the interplay with ppGpp and DksA (65). Interestingly, two relatively small proteins (<20 kDa) with unknown function, YajQ und YhfA, were upregulated in several of replicates. *yajQ* was described as “survival gene” under cell death conditions and was shown to increase resistance against hydrogen peroxide (66, 67). YfgM, yet another stress protein induced in several replicates after exposure to the inhibitors, is a bifunctional inner-membrane protein. It is involved in the periplasmic chaperone network and cytoplasmic stress adaptation (68). Like LpxC, it is a substrate of the FtsH protease and prone to degradation in a growth phase-dependent manner. Since *yfgM* is not transcriptionally induced in response to LpxC inhibition (32, 69), its increased abundance suggests a stabilization against FtsH-mediated turnover.

Compound BB-78485 deviated from the other four inhibitors in several respects. MIC and PEC concentrations were much higher, and the cellular responses were different. Notably, this compound was significantly less potent against *P. aeruginosa,* but had an increased activity in a semipermeable *P. aeruginosa* strain (22). As the determined inhibitory constant of BB-78485 (K_i_= 20 nM) is in between those determined for L-161,240 (K_i_= 50 nM) and CHIR-090 (K_i_= 4 nM) (30), the observed differences could be related to a lower uptake. This, however, cannot account for substantially different physiological responses to BB-78485. It did not elicit LPE formation but induced more than 40 specific marker proteins. The higher concentration used might have induced off-target binding resulting in adverse effects. Moreover, the low incorporation of L-[^35^S] methionine suggests some translation defects. These findings suggest that BB-78485 binds to other targets apart from LpxC. BB-78485 is a sulfonamide derivative that inhibits folic acid synthesis. This is an essential metabolic pathway for the synthesis of new DNA building blocks such as purines and pyrimidines (70). Since several DNA synthesis-relevant proteins such as PurC, PyrB, PyrF and PyrG are upregulated almost exclusively upon BB-78485 exposure, this could indicate a dual-mode of action for this compound, which could be beneficial for medical applications.

### 3.2. Species-specific responses to LpxC inhibitors in Gram-negative bacteria

While some LpxC inhibitors such as CHIR-090 have potent efficacy against *E. coli* and *P. aeruginosa* comparable to that of ciprofloxacin (71), they are less effective against other bacteria (72). LpxC is conserved among Gram-negatives, but the sequences are quite divergent. The *E. coli* protein shares 98% homology to LpxC from *S. enterica* and 57% to that of *P. aeruginosa*, but only shares 30% similarity to the *Aquifex aeolicus* protein (73). The species-specific efficacy of the inhibitors is probably related to the different properties of the active binding site. While *Ec*LpxC has a relatively spacious active site that can accommodate sterically bulky compounds such as BB-78485, this sulfonamide fails to bind to the smaller binding cavity of *Aa*LpxC (74).

In comparison to *E. coli*, the inhibitor CHIR-090 elicits a substantially different cellular response in *P. aeruginosa* (45). Instead of LpxC or FAB enzymes, upregulated marker proteins included proteins involved in precursor supply, iron acquisition, DNA replication and export systems. As in *E. coli*, DksA and ROS detoxification were among the induced proteins. The overall proteome pattern suggested that multiple cellular processes were perturbed by CHIR-090 treatment in *P. aeruginosa*.

The major reason for the difference in the response between *E. coli* and *P. aeruginosa* presumably lies in the entirely different mode of regulation of LPS biosynthesis. While the LpxC proteins of *E. coli*, *S. enterica* and *Yersinia pseudotuberculosis* are prone to FtsH-mediated proteolysis, this is not the case for *P. aeruginosa* and *A. aeolicus* LpxC, which are stable proteins (73). Instead of the tight coupling between LPS and PL biosynthesis in *E. coli* (Fig. 10), coordination of OM biosynthesis in *P. aeruginosa* is coordinated with cell wall biosynthesis. It was recently shown that MurA, the first enzyme in the peptidoglycan biosynthesis pathway, interacts with and influences the activity of LpxC (75). This proteolysis-independent strategy ensures a synchronized synthesis of cell envelope components. Consistent with that finding, CHIR-090 treatment in *P. aeruginosa* induced GlmU, which provides UDP-GlcNAc, the common precursor of LPS and peptidoglycan synthesis (45). This does not exclude a regulatory interconnection between LPS and PL biosynthesis in *P. aeruginosa*. One piece of evidence for this is actually the appearance of suppressor mutation in *fabG* in CHIR-090-resistant strains strongly suggesting a link to fatty acid biosynthesis (46). Such species-specific responses can be observed not only for LpxC inhibitors but also for other antibiotics such as allicin or rifampicin (45, 76), indicating a different “wiring” of the regulatory network in Gram-negative bacteria.

### 3.3. Essentiality as a weak point - LPS synthesis offers many promising targets

LPS biosynthesis is a complex, multi-factorial process involving a large number of enzymes and regulatory proteins (Fig. 10). Therefore, it is conceivable that essential proteins other than LpxC in this pathway are equally suitable drug targets. Indeed, inhibitors of the enzymes LpxA, LpxD, LpxH, LpxK, the ABC transporter MsbA, and the Lpt complex have been reported (77–82). Aside from evaluating their antimicrobial potency and toxicity to mammalian cells, it will be important to carefully assess their potential to develop suppressor mutations as it has recently been evaluated for LpxC and the two inhibitors CHIR-090 and PF-04753299 (16). An invaluable advantage of LPS inhibitors stems from the fact that LPS, PL and cell wall biosynthesis are three essential and tightly interlinked processes (18, 19, 83). It is easily conceivable that interference with one process may have far-reaching effects into the overall cell envelope homeostasis. Antibiotic combination therapy targeting two of these crucial pathways may therefore be a viable approach, in particular for combating multidrug-resistant bacteria (84, 85). Administration of drug combinations can circumvent the formation of resistant subpopulations as it was shown for *P. aeruginosa* with CHIR-090 and colistin (86). Here, we showed that pre-treatment of cells with low doses of LpxC inhibitors increased the susceptibility of *E. coli* to vancomycin but not to ampicillin. Both antibiotics interfere with peptidoglycan synthesis. Ampicillin reaches the cell wall of both Gram-negative and positive cells and some resistance is achieved by the constitutive production of β-lactamases in *E. coli* (87). In contrast, vancomycin is used exclusively to treat Gram-positive infections (88, 89) because Gram-negative bacteria like *E. coli* are intrinsically resistant to it due to the permeability barrier formed by the outer membrane. Hence, vancomycin can only harm Gram-negative cells with compromised OMs, e.g. an LPS-lacking *Acinetobacter baumannii* strain (90). Therapy of drug-resistant Gram-negative pathogens by a combination of LpxC inhibitors and vancomycin may thus have the potential to cure poorly treatable infections.

## 4. Material and methods

The utilized LpxC inhibitors CHIR-090 (Axon MedChem), L-161,240 (AdooQ), BB-78485 (Aobius), PF-04753299 (Sigma-Aldrich), PF-05081090 (Axon MedChem) were purchased online. The LpxC inhibitors were dissolved in DMSO at a stock solution of 5 mg/ml.

**Strains and growth media**. The *lon*-deficient *E. coli* BL21 [DE3] (91) strain was used for protein overproduction. Therefore, the strain harbored a high-copy expression vector and was cultivated in Luria-Bertani (LB) medium (92) supplemented with appropriate antibiotics at 37°C and 180 rpm. *E. coli* BW25113 and the two Keio-library derived mutants Δ*pldA* and Δ*pagP* (36) as well as *E. coli* W3110 (93) and the efflux pump-lacking mutant *E. coli* W3110 Δ*tolC* were cultivated at 37°C in LB or in M9 minimal medium (8.09 mM Na_2_HPO_4_, 12.79 mM KH_2_PO_4_, 8.56 mM NaCl, 18.69 mM NH_4_Cl, 22 mM glucose, 100 μM CaCl_2_, 2 mM MgSO_4_, 29.65 μM thiaminium dichloride, 46.26 μM H_3_BO_3_, 0.01 μM MnCl_2_, 0.77 μM ZnSO_4_, 1.61 μM Na_2_MoO_4_, 0.49 μM CuSO_4_, 0.17 μM Co(NO_3_)_2_).

### Protein production and purification

1. *E. coli* BL21 [DE3] harbored the plasmid pBO2382 (pASKiba5+ derivative), which encoded for N-terminal His_6_-tagged LpxC (73). Cells were cultivated until the exponential phase (OD_600nm_ of 0.5) and for the protein overproduction, the culture was shifted to 30°C and 20 ng/ml AHT was added. After 3 hours, cells were harvested, pellets were washed twice with 20 mM HEPES and 150 mM NaCl (pH 8) and then stored at −20°C. For cell disruption, the pellets were thawed on ice and resuspended in buffer (20 mM HEPES, 150 mM NaCl, spatula tip lysozyme, DNase, and RNase and 1x cOmplete EDTA-free protease inhibitor, pH 8). Lysis was performed three times at 900 PSI with the FrenchPress system (ThermoElectron). Then, cell debris and supernatant were separated via centrifugation for 20 min at 4000 x *g* at 4°C. Finally, the supernatant was load onto equilibrated purification columns (20mM HEPES, 150 mM NaCl) with NiNTA (nitrilotriacetic acid) and incubated for 1h. The NiNTA resin was washed with 10 ml buffer 2 (20 mM HEPES (pH 8.0), 500 mM NaCl, 10 % (w/v) glycerol, 50 mM imidazole), buffer 3 (20 mM HEPES (pH 8.0), 300 mM NaCl, 10% (w/v) glycerol, 50 mM imidazole) and buffer 4 (20 mM HEPES (pH 8.0), 150 mM NaCl, 10% (w/v) glycerol, 50 mM imidazole) and proteins of interest were eluted with buffer 5 (20 mM HEPES (pH 8.0), 150 mM NaCl, 10 % (w/v) glycerol, 250 mM imidazole). The protein elution buffer was exchanged using Amicon® Ultra-4 Centrifugal Filter Units with a size exclusion of either 10 to 20 mM Hepes and 150 mM NaCl (pH 8). Final protein concentration was determined with a Bradford assay (94).

### SDS-PAGE analysis and immunodetection

For the analysis of the *in vivo* degradation experiments or the detection of the LpxC abundance, the harvested cells were resuspended according to their optical density (100 µl for OD_580nm_= 1) in TE buffer (10 mM Tris/HCl, pH 8; 1 mM EDTA) and 5 x protein loading dye (2% (w/v) SDS, 0.1% (w/v) bromophenol blue, 10% glycerol, 50 mM Tris/HCl, pH 6.8). All samples were heated for 10 min at 100°C, centrifuged for 1 min at 16.000 x *g* and then 15 µl were loaded on a 12% SDS-PAGE, followed by a western transfer using standard protocols (92). The nitrocellulose membranes were blocked with 3% BSA-TBST for 1 h followed by three washing steps with TBST for 10 min. The blocked membrane was incubated with a polyclonal rabbit anti-LpxC antibody (diluted 1:20000 in TBST) for 60 min, followed by three-times washing with TBST for 5 min and subsequent incubation with the goat anti-rabbit IgG (H+L)-HRP conjugate (BioRad) (diluted 1:3000 in TBST) for again 60 min. Finally, the membranes were washed three times with TBST for 10 min and proteins were detected via chemiluminescence using Immobilon® Forte Western HRP substrate (Merck). For the multiplex detection of native LpxC and the LPS level 12% stain-free FastCast Acrylamide gels (BioRad) were prepared according to manual instructions. Samples were prepared and loaded as described before. Prior to blotting, the overall protein was visualized with the stain-free channel of the ChemiDoc MP BioRad Imager. Proteins were then transferred onto nitrocellulose using the Trans-Blot® Turbo™ Transfer System (BioRad) with the recommended manufacturers consumables. After 5 min of blocking in EveryBlot blocking buffer (BioRad), a mouse anti-LPS core monoclonal antibody mAb WN1 222-5 (Hycultbiotech) (1:4000) and the polyclonal rabbit anti-LpxC antibody (1:20000) were added and incubated for 60 min at RT. The membrane was washed three times with TBST for 5 min and the blots were then incubated for 60 min with the second antibodies, StarBright Blue 700 goat anti-mouse IgG (BioRad) and StarBright Blue 520 goat anti-rabbit lgG (BioRad), diluted 1:2500 in EveryBlot blocking buffer. After three times of washing with TBST, the membranes were rinsed with TBS, then dried, and the fluorescence signal was detected with the ChemiDoc MP BioRad imager. Quantification of total protein and specific signals were analyzed using the ImageLab software (BioRad).

### Thermal shift assay to verify LpxC inhibitor binding

20 µl of freshly purified His_6_-tagged LpxC (supplemented with 12.7 µM DTT, final assay conc.: 12.5 µM) were mixed with 3 µl BODIPY^TM^ FL L-Cystine (final assay conc.: 2 µM) and 2 µl of protein buffer, DMSO or the LpxC inhibitors (final assay conc.: 50 µM) in a clear 96 well plate. The plate was sealed with a transparent foil and centrifuged for 1 min at 2000x*g*. The fluorescence was measured at 512 nm along a temperature gradient from 20 to 100 °C (0.5°C increase every 10 s) in the CFX Connect^TM^ real-time thermocycler (BioRad) with the Cfx Maestro^TM^ software. Each measurement was performed in technical triplicates.

### DiSC3 Assay to analyze the membrane depolarization

Exponential *E. coli* W3110 cells were adjusted with LB in a volume of 1 ml to an OD_580nm_ of 0.5. Cells were washed twice with 1 ml DiSC3 buffer (5 mM HEPES pH 8, 5 mM glucose, 0.2 mM EDTA). 90 µl of the cell suspension was transferred into a flat-black 96 well plate and immediately before starting the measurement, 1 µl of the fluorescent, membrane-integrative probe DiSC3 (stock: 50 µM) was added. Fluorescence was measured in the Tecan infinite M PLEX plate reader till the signal stagnated (excitation: 622 nm; emission: 670 nm; temperature: 18°C; intervals of 20 s). Then, the plate was taken out briefly and 10 µl of DMSO, polymyxin B or LpxC inhibitors (CHIR-090, L-161,240, BB-78485, PF-04753299 and PF-05081090) (final conc.: 16 µg/ml) were added per well. The fluorescence was measured for 7 min with the previous settings. The experiment was performed in biological triplicates and each time with three technical replicates. First, the mean value of the technical replicates within one experiment were calculated. The values obtained directly before addition of the compounds were normalized to 1 and the other values were adjusted accordingly. Then, the normalized average of the biological replicates and the standard deviation were calculated. These values were used for the graph generation.

### Determination of the minimal inhibitory concentration (MIC)

Exponential cells of 1. *E. coli* W3110 and the efflux pump deficient *E. coli* W3110 Δ*tolC* were used for determination of the MIC for the LpxC inhibitors in M9 minimal medium. Cell density was adjusted to 5 x 10^5^ cells/ml (OD_580nm_ of 1 ≙ 6 x 10^7^ cells/ml) and 2 ml were transferred to test tubes. The growth inhibiting effect of the inhibitors were tested in a dilution assay (0, 0.001, 0.005, 0.01, 0.025, 0.05, 0.1, 0.2, 0.3 and if necessary, also 0.5, 0.75, 1, 2.5, 5, 7.5, 10 and 15 µg/ml). After 18 hours of incubation at 37°C in a tube rotor, growth inhibition was determined by measuring the OD_580nm_ of the test tubes. The concentration of the sample in which no growth was observed, was determined as the MIC for the corresponding compound.

### **Determination of the physiologically effective** concentration **(PEC).**

The PEC is defined as the concentration required to inhibit growth within the first two hours after addition of a compound by a minimum of 30%. Exponential *E. coli* W3110 cells grown in M9 minimal media were treated with different concentrations of the LpxC inhibitors depending on the previously determined MICs (¾ MIC, ½ MIC, ¼ MIC). The OD_580nm_ was measured every hour for five hours after compound addition.

### **Microscopy of LpxC inhibitor treated** cells

A main culture of exponential *E. coli* W3110 cells (OD_580nm_ of 0.5) was divided into six subcultures of 5 ml each. The cultures were treated with 1.5 x MIC of the LpxC inhibitors (Table 1) or with DMSO (final concentration 0.25%) as negative control. After five hours, cell density was adjusted to an OD_580nm_ of 0.5. Perforated cells were stained with 30 µg/ml propidium iodide (PI), a DNA dye that cannot pass intact but damaged membranes. 5 µl of the cells were applied to a slide with an agar patch (1 x PBS, 1.5% agarose). The cells were examined under 100x magnification with an Olympus fluorescence microscope. Images were taken in brightfield to obtain all cells and in the TxRed channel to see the PI-stained, i.e. damaged cells. To statistically evaluate the effect of the inhibitors on the cell shape, the cell length and width was measured with the ImageJ software from at least 100 cells per treatment. The percentage viability of the cells was determined after five hours by subtracting the ratio of cells stained with PI to cells counted in brightfield (n>100) from 1 and multiplying the final value by 100. The viability of the cells before addition of the compounds was set at 100 % (t= 0h).

### Survival spotting assay.

Exponential *E. coli* W3110 cells, grown in LB, were equilibrated to OD_580nm_ of 0.5 and were then treated with ¼ MIC of the LpxC inhibitors, or as controls with DMSO or 300 µg/ml spectinomycin, for two hours. The volume required for a final cell density of OD_580nm_ 0.5 in 1 ml was harvested. To remove the compounds, cells were washed once with 0.9% NaCl and the cell pellet was finally resuspended in 1 ml 0.9% NaCl. Starting from this cell suspension, a dilution series was prepared with a factor of 10. 3 µl of each dilution were then spotted onto LB, LB with ampicillin (2 µg/ml), LB with 1% SDS and 0.5 mM EDTA or LB with vancomycin (150 µg/ml) and incubated overnight at 37 °C. The next day, colonies of one dilution level were counted to determine the cell viability to a specific treatment after incubation with LpxC inhibitors. The experiment was performed in triplicates.

### **Lipid analysis by one-dimensional thin layer** chromatography **(1D TLC)**

Exponential *E. coli* W3110 or *E. coli* BW25113 and the corresponding mutants Δ*pldA* and Δ*pagP*, grown in M9 minimal medium, were supplemented with ½ MIC of the LpxC inhibitors or as control with DMSO for 2 hours. Then, the OD_580nm_ was measured and the required volume for 1 ml cell suspension with an OD_580nm_ of 10 for each culture was calculated. After harvesting by centrifugation, the cell pellets were resuspended in 100 µl distilled water and lipids were isolated as described before (95). The organic phase was transferred in a new tube and the liquid was evaporated in a speed vacuum centrifuge. Finally, the lipids were resuspended in methanol:chloroform (1:1) and subjected to thin layer chromatography (TLC) using silica gel 60 plates (Merck). Total lipids were separated using chloroform:methanol:water (65:25:4) as mobile phase and phospholipids were visualized using molybdenum blue spray reagent(96). Lipids with free amino groups were stained with a ninhydrin staining solution (1.5% (w/v) ninhydrin and 3% (v/v) acetic acid in n-butanol) and visualized by incubation at 140°C for 10 min. Lipids were identified by comparison of the retention behavior of commercially available PL. The experiment was performed in biological triplicates.

### **Preparation of cytoplasmic L-[^35^S] methionine-**labeled **protein extracts.**

A culture of *E. coli* W3110 (OD_580nm_ of 0.35) grown in M9 minimal medium at 37 °C, was split into 20 ml subcultures. The growth after addition of the PEC (Fig. 4A) of the respective LpxC inhibitor was followed by measuring the OD_580nm_ every hour. For radioactive pulse-labeling, 10 min after the addition of the LpxC inhibitor, 5 ml of the subcultures were transferred into a new flask and mixed with 1.8 MBq L-[^35^S]-methionine (Hartmann analytic). After 5 min, the methionine incorporation was blocked by addition of 1 mg/ml chloramphenicol, an excess of nonradioactive L-methionine (10 mM) and by direct transfer of the cultures on ice. After cell harvest, pellets were washed three times (100 mM Tris pH 7.5, 1 mM EDTA) and for disruption via sonication in the VialTweeter (Hielscher) cells were resuspended in 10 mM Tris buffer (pH 7.5) containing 1.39 mM phenylmethylsulfonylfluoride as protease inhibitor. The debris and membrane-bound proteins were separated from the soluble protein fraction by centrifugation at 13.000 x *g* for 20 min. For the determination of the protein biosynthesis rates, 1 µl of each soluble protein fraction was spotted onto Whatman paper. Then, proteins were precipitated by incubation in 20% trichloroacetic acid, followed by 10% trichloroacetic acid and then 96% ethanol. After drying the Whatman papers at room temperature, the incorporation rates of radioactive L-[^35^S]-methionine were measured with a scintillation counter. Protein concentration was determined using a Bradford-based Roti NanoQuant assay (Roth). The volume corresponding to 55 µg (radioactive) or 300 µg (non-radioactive) of total protein was transferred in a separate tube and evaporated in a SpeedVac centrifuge.

### 2D-PAGE and identification of regulated proteins by mass spectrometry (MS).

Analytical and preparative 2D-PAGE were performed as described earlier (97). In short, for the radioactive gels 55 µg and for the non-radioactive gels 300 µg lyophilized protein were resuspended in 500 µl rehydration solution (7 M urea, 2 M thiourea, 65 mM CHAPS (3-[(3-cholamidopropyl)dimethylammonio]-1-propanesulfonate), 0.5 % (v/v) Triton X-100, 1.04% PharmalyteTM 3 – 10 (GE Healthcare), 50 mM DTT and trace amounts of bromophenol blue), of which 450 µl were subjected to isoelectric focusing on pH 4-7 IPG strips (GE Healthcare), followed by SDS-PAGE with 12 % acrylamide gels. Pulse-labeled proteins were visualized by exposing the dried gels to phosphoscreens for several hours up to days prior to detection with the Typhoon Trio+ Imager (GE Healthcare). Regulated proteins were identified using the Delta2D software (Decodon) as described before (98). The threshold for “regulated spots” was at least 1.6-fold upregulation in three independent biological replicates.

The total protein of the non-radioactive gels was stained with RuBPS and were used to excise the identified “regulated protein spots” for MS analysis. After tryptic in-gel digestion (99), 5 µl were injected onto an ACQUITY UPLC I-Class System (Waters) equipped with an ACQUITY UPLC BEH C_18_ column at 40°C (Waters, particle size 1.7 µm, column dimensions: 2.1 x 50 mm). A 7 min gradient with H_2_O and acetonitrile (ACN), each with 0.1% formic acid (FA), was used with a flow rate of 0.4 ml/min (Table S2). Masses eluted before 0.5 min and after 5 min were not injected into the mass spectrometer.

Data-independent continuous MS^E^ measurements were performed with a Vion IMS QToF (Waters) with an ESI source in positive sensitivity mode. Masses in a range of 50 to 2000 m/z were detected with 0.2 s per scan and leucine enkephalin being injected as a reference mass at the beginning of every sample and after 5 min. Used parameters: capillary voltage 0.8 kV, sample cone voltage 40 V, source offset voltage 80 V, cone gas flow 50 l/h, desolvation gas flow 1000 l/h, source temperature 150 °C, desolvation temperature 550 °C, collision gas N_2_, collision low energy 6 V, collision high energy ramp 28-60 V.

Recorded data was exported to waters.raw format using UNIFI (Waters, version 1.9.13) and processed with ProteinLynx Global Server (Waters, version 3.0.3) as described previously (76) with modified energy thresholds: low energy 50 counts, high energy 10 counts. A databank containing 4212 proteins for *E. coli* W3110 (NCBI reference sequence NC_007779.1) was used, following the method described in Wüllner et al., 2019.

### ***In vivo* degradation experiment to** study **LpxC stability.**

1. *E. coli* W3110 cells were cultivated in LB in a water bath shaker until stationary phase (OD_580nm_ of 3). Prior to addition of the inhibitors, 1 ml of sample was taken (−15) and then the culture was spiked with the respective MIC of the compounds (Table 1). After 15 min, 200 µg/ml chloramphenicol was added to inhibit translation and 1 ml samples were taken after 0, 5, 10, 20, 30, 60, 90 and 120 min. All samples were directly frozen in liquid nitrogen. The native LpxC level of each culture was analyzed immunologically by at least by three biological replicates.

## Data availability

The mass spectrometry proteomics data have been deposited to the ProteomeXchange Consortium via the PRIDE (100) partner repository with the dataset identifier PXD043195 (Reviewers can access the data using the reviewer account: Username: password: kTSHTDMC

## Acknowledgments

We thank Pascal Dietze for help with 2D gel electrophoresis and RUBION for support. This study was funded by the German Research Foundation (DFG; Research Training Group 2341 “Microbial Substrate Conversion (MiCon)” and the Priority Program SPP1879 “Nucleotide second messenger signaling in bacteria”), the German Federal State of North Rhine-Westphalia and the European Union, European Regional Development Fund, Investing in your future (Research Infrastructure “Center for System-based Antibiotic Research (CESAR)”) to JEB. Figures were created with BioRender.com.

The authors declare that they have no conflicts of interest. All data generated or analyzed during this study are included in this published article (and its supplementary information files).

